# Interfacing Spinal Motor Units in Non-Human Primates Identifies a Principal Neural Component for Force Control Constrained by the Size Principle

**DOI:** 10.1101/2021.12.07.471592

**Authors:** Alessandro Del Vecchio, Rachael H. A. Jones, Ian S. Schofield, Thomas M Kinfe, Jaime Ibáñez, Dario Farina, Stuart N. Baker

## Abstract

Motor units convert the last neural code of movement into muscle forces. The classic view of motor unit control is that the central nervous system sends common synaptic inputs to motoneuron pools and that motoneurons respond in an orderly fashion dictated by the size principle. This view however is in contrast with the large number of dimensions observed in motor cortex which may allow individual and flexible control of motor units. Evidence for flexible control of motor units may be obtained by tracking motor units longitudinally during the performance of tasks with some level of behavioural variability. Here we identified and tracked populations of motor units in the brachioradialis muscle of two macaque monkeys during ten sessions spanning over one month during high force isometric contractions with a broad range of rate of force development (1.8 – 38.6 N·m·s^-1^). During the same sessions we recorded intramuscular EMG signals from 16 arm muscles of both limbs and elicited the full recruitment through neural stimulation of the median and deep radial nerves. We found a very stable recruitment order and discharge characteristics of the motor units over sessions and contraction trials. The small deviations from orderly recruitment were observed between motor units with close recruitment thresholds, and only during high rate of force development. Moreover, we also found that one component explained more than ~50% of the motor unit discharge rate variance, and that the remaining components could be described as a time-shifted version of the first, as it could be predicted from the interplay between the size principle of recruitment and one common input. In conclusion, our results show that motoneurons recruitment is determined by the interplay of the size principle and common input and that this recruitment scheme is not violated over time nor by the speed of the contractions.

## INTRODUCTION

Theories of motor control are grounded on recording spinal motor unit activity during voluntary force contractions (*1–4*). Accurate understanding of motor unit function reveals in a direct way the strategies used by the nervous system to control and coordinate muscle forces (*4*). Generation of force is believed to occur by a combination of recruitment and rate coding of spinal motor neurons. While it is often assumed that recruitment order and rate coding are determined by the size principle (*5, 6*) and the common inputs that the motor neurons in a pool receive (*2*), some studies have challenged this view by proposing a more flexible motor unit control (*7–9*). Although previous evidence supports the size principle during isometric contractions (*10, 11*), these results have been challenged by the possibility that the motor cortex could provide independent input to spinal motoneurons. Moreover, it is still unclear if the high correlations in motor unit output (*2, 12-14*) have a functional origin or represent a physiological epiphenomenon.

The current lack of definitive evidence for size principle and common input during recruitment with force modulation is due to technical limitations. Accurate measures of the recruitment order and common input necessitate multiple recordings from as many units as possible and the tracking of the same motor units across different days and across rates of muscle force development (*4, 7, 8, 10, 11, 15-18*). Currently, no studies tracked the same population of motor units in longitudinal experiments during natural tasks in non-human primates. Such tracking of the same population of neurons is crucial to infer functional behaviour. This is even more important when testing intrinsic properties of motoneurons, such as those associated with the size principle. One way to identify motor unit activity during natural tasks is to insert percutaneous wire electrodes into muscles. However, these electrodes may yield limited signal quality and limited number of detected motor units.

By tracking the behaviour of the same motor neurons across multiple experimental sessions with a new non-invasive neural interface consisting of high-density grids of electrodes placed on the muscle, we investigated for the first time the variability in motoneuron recruitment and discharge characteristics over a period of one month in two monkeys during natural contractions. The tracking of a relatively large population of spinal motor units during contractions at different rates of force development allowed us to define the neural strategies accomplished by the central nervous system to control muscle force. Moreover, it was possible to investigate the associations between recruitment of motoneurons and estimates of common synaptic inputs.

We found a very small day-to-day and trial-to-trial variability in recruitment order and rate coding, suggesting consistent control of the population of motoneuron ensembles. Moreover, with a factorization method we demonstrated that one common input component was sufficient to explain motor unit recruitment. The application of this approach in a primate species with a motor system closely similar to humans opens the future possibility of combining multiple single motor unit measurements with invasive recordings from central pathways. This has the potential to yield substantial new insights into the anatomical source of common drive during different motor tasks.

## RESULTS

### Motor unit decomposition and tracking

We describe the strategies of control of macaque motor units and evaluate the performance of a new non-invasive neural interface framework to monitor the changes in the number and properties of longitudinally tracked units over 10 experimental days (gathered over one month) in two animals.

We decomposed spike trains of individual motor units from high-density EMG signals using blind source separation techniques (Figure 1A; see details in Methods). After this process, the spike trains belonging to each decomposed motor unit were used to estimate the average 2D waveform of the corresponding action potentials (Figure 1A shows one column of the recording grid). The motor unit waveforms were used to track the same motor unit with a 2D cross-correlation function (*19, 20*). Figure 1B shows the raster plot of all motor units across the ten days for Monkey MI. The y-axis in Figure 1B shows the total number of identified motoneurons across days (color-coded). The central panel of Figure 1B shows an example of force signal and raster plot of the motor units during a contraction.

**Figure 1.**
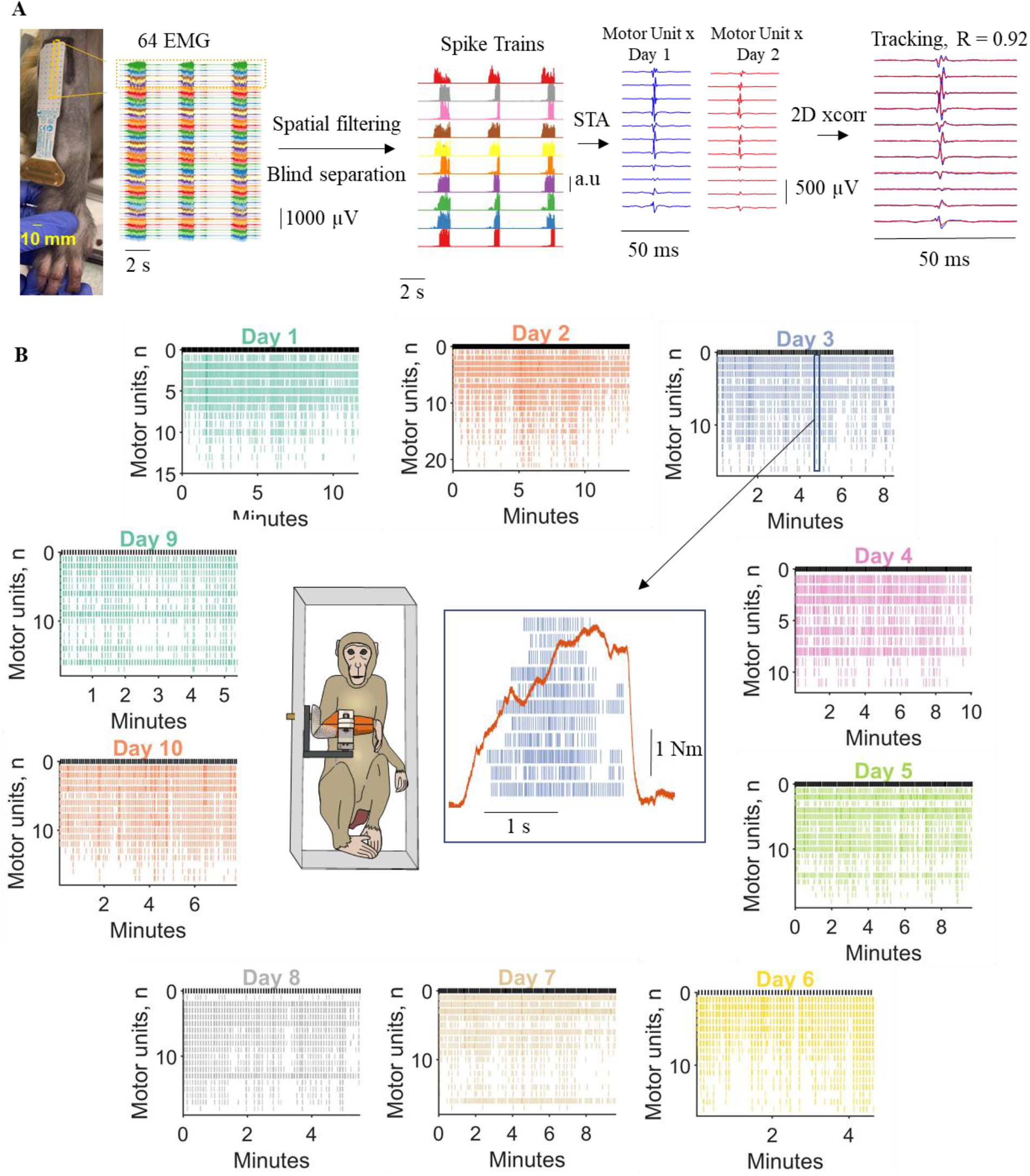
Motor unit decomposition in awake behaving macaques, experimental framework and analysis. **A.** From left to right, sixty-four monopolar EMG signals during three individual contractions. Each contraction lasted approximately 2 seconds. The monopolar EMG signals were spatially filtered with a double-differential derivation. After this process, blind source separation identified the spike trains belonging to individual motor units. The spike trains for each motor unit were used to spike trigger the average 2D motor unit waveform. The 2D motor unit waveforms were used for the longitudinal tracking, through a 2D cross correlation function. **B.** Monkey 1 (MI) individual motoneuron spike trains across the 10 days (colour coded). Note that during the different days we identified a relatively similar number of motor units. The centre of the figure shows the experimental setup and an individual voluntary contraction (force signal in red) extracted from Day 3. *STA = Spike-triggered average.

On average, each recording session (one per day, ten days in total) lasted 9.8 ± 2.5 min (Monkey MI) and 8.6 ± 2.8 min (Monkey MA). During these sessions, the monkeys performed on average 118.0 ± 30.1 (MI) and 103.5 ± 33.9 (MA) contractions, that were used for the subsequent EMG analyses. The monkeys were instructed to reach a target without a specific training on the rate of force development. Therefore, we obtained a relatively large variance in rate of force development and motor unit recruitment speeds across contractions. During these contractions the rate of force development ranged widely, with an average and standard deviation of 6.44 ± 4.00 N·m·s^-1^ (range 1.86 – 38.66 N·m·s^-1^). Moreover, the peak force obtained across days also showed high variability, spanning two-fold maximum EMG amplitudes.

We identified a total of 389 motor units (192 MI and 197 for MA) in the individual recordings. Of these, only a subset (Figure 2) could be tracked and reliably matched with a unit from one or more different days on the basis of a two-dimensional correlation coefficients R>0.7 (see details in Methods). The average number of identified motor units for each experimental session was 19.2 ± 2.97 and 19.7 ± 2.4 (mean and standard deviation), for MI and MA respectively. We were able to track on average 9.07 ± 1.06 and 8.13 ± 2.08 motor units across all 10 days. Figure 2 shows the total number of identified motor units at each day and the number of tracked motor units across sessions, for the two monkeys. The upper panel of Figure 2 shows examples of 2D and 3D motor unit waveforms as well as the total number of motor units across contractions and days (bottom panels F-I). Figure 2F-I depicts the total number of motor units decomposed on each day for both monkeys. The right panels (Fig. 2G-I) show the individual motor units that were tracked across the different days (all possible combinations). Note that the largest number of units in these bar plots correspond to the units recorded during the examined day, which are highlighted with a black edged bar (Fig 2G and 2I). The number of the tracked units across days was lower than the total number of identified motoneurons (on average 19.45 vs. 8.60) because small changes in the proportion of recruited motor units challenge the tracking procedure. We previously obtained a very similar result in humans (*20*) due to different target forces and day-to-day variability.

**Figure 2.**
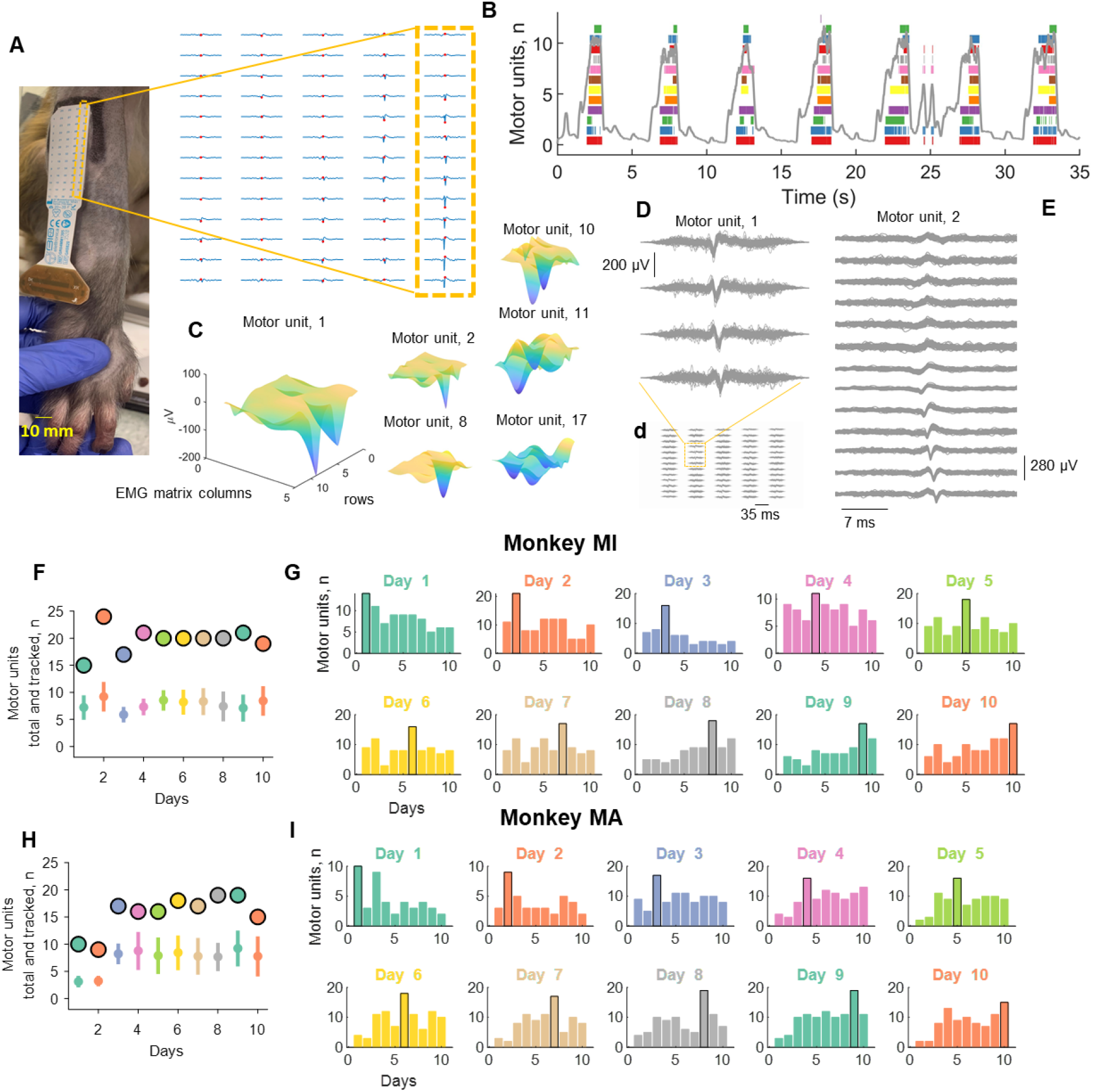
Motor unit action potentials and total numbers of identified and tracked motor units across the 10 days (color-coded). **A**. Two-dimensional motor unit action potential propagating under the high-density EMG electrode array. The highlighted yellow inset shows the respective column and row of the high-density EMG matrix during the experiment. **B.** Raster plot of 12 identified motor units (color-coded) for seven representative contractions. **C**. Three-dimensional representation of the motor unit action potential in a specific time instant (highlighted with a red dot in A). Note that each action potential has a unique 3D signature which allows the independent component analysis to converge to the time-series of discharge timings of the motor unit. **D**. Shimmer plots for two action potential waveforms. Each action potential was averaged across an individual contraction and then superimposed across all contractions for a specific day. Note the high similarity across channels for two representative motor units. **F.** The total number of identified motor units across the 10 days (black edge circles) and tracked motor units (open circles with vertical line depicting the standard deviation) for monkey MI and for MA (**H**). **G-I.** Bar plot of the number of motor units that were successfully tracked across the 10 days (color-coded). Note that the black edged bar plot corresponds to the number of motor units that were identified at the respective day and used for tracking those motor units in the other days.

Despite the number of tracked motor units being lower than the number of identified motor units, the discharge characteristics of the tracked motor units was highly correlated across sessions over the full duration of the experiments (~1 month), as described in the following section.

### Motor unit identification validity

The motor unit action potential similarity across sessions was assessed with the two-dimensional (2D) cross-correlation function (see details in Methods). Because the motor unit action potential waveform and motor unit discharge characteristics are independent, we first computed quality measures of decomposition based on the action potential waveform, and successively we computed correlation measures between the tracked motor units firing characteristics (discharge rate and recruitment threshold across days).

The consistency of each motor unit action potential that was accepted to belong to the same cluster, was very high (Silhouette measure averaged across all the identified motor units and the 10 days, 0.91 ± 0.01 and 0.92 ± 0.01, for MI and MA respectively). Silhouette measures above 0.9 have been associated with highly accurate decomposition with respect to intramuscular EMG signals (*21*). Moreover, the tracked units across sessions exhibited very high 2D correlation coefficients of the motor unit waveform (>0.7 for the tracked units) and similar discharge rates across the different days. Figure 2D shows the action potentials that were spike-trigger averaged across the individual contractions (all the action potentials for a representative contraction were used to generate the motor unit action potential waveform, across all 64 channels). The variability in the action potential waveforms across contractions for the same day were minimal, with action potentials 2D correlation values always above 0.9. This indicates very high reliability in identifying the same motor unit across contractions.

### Physiological characteristics of macaque motor units

Figure 3 shows the discharge characteristics of the tracked motor units across and between days. The inter-day motor unit discharge rate variability was very low, at 3.51 and 5.41 % for MI and MA respectively. For Monkey MI, the bivariate Pearson correlation coefficients between the average discharge rate across the different days were significant in all cases (P<0.001 after Bonferroni correction, Fig.3A-B). Indeed, the absolute differences in discharge rate across the units over the different days (Fig. 3C) was very low (0.14 ± 3.45 spikes/s). For Monkey MA, the results were similar, although with a smaller number of reliably decomposed motor units during the first two days (Fig. 2B), that resulted in poorer tracking performance during those two days (Fig 3. D-E). However, the lower number of motor units did not change the performance of the tracking algorithm and discharge characteristics of the units. There was a very small variability in discharge rate of the tracked units and corresponded to 0.09 ± 3.12 spikes/s, with an average discharge rate across the ten days for all the identified motor units of 41.77 ± 1.46 and 38.42 ± 2.07 (spikes/s), for MI and MA respectively.

**Figure 3.**
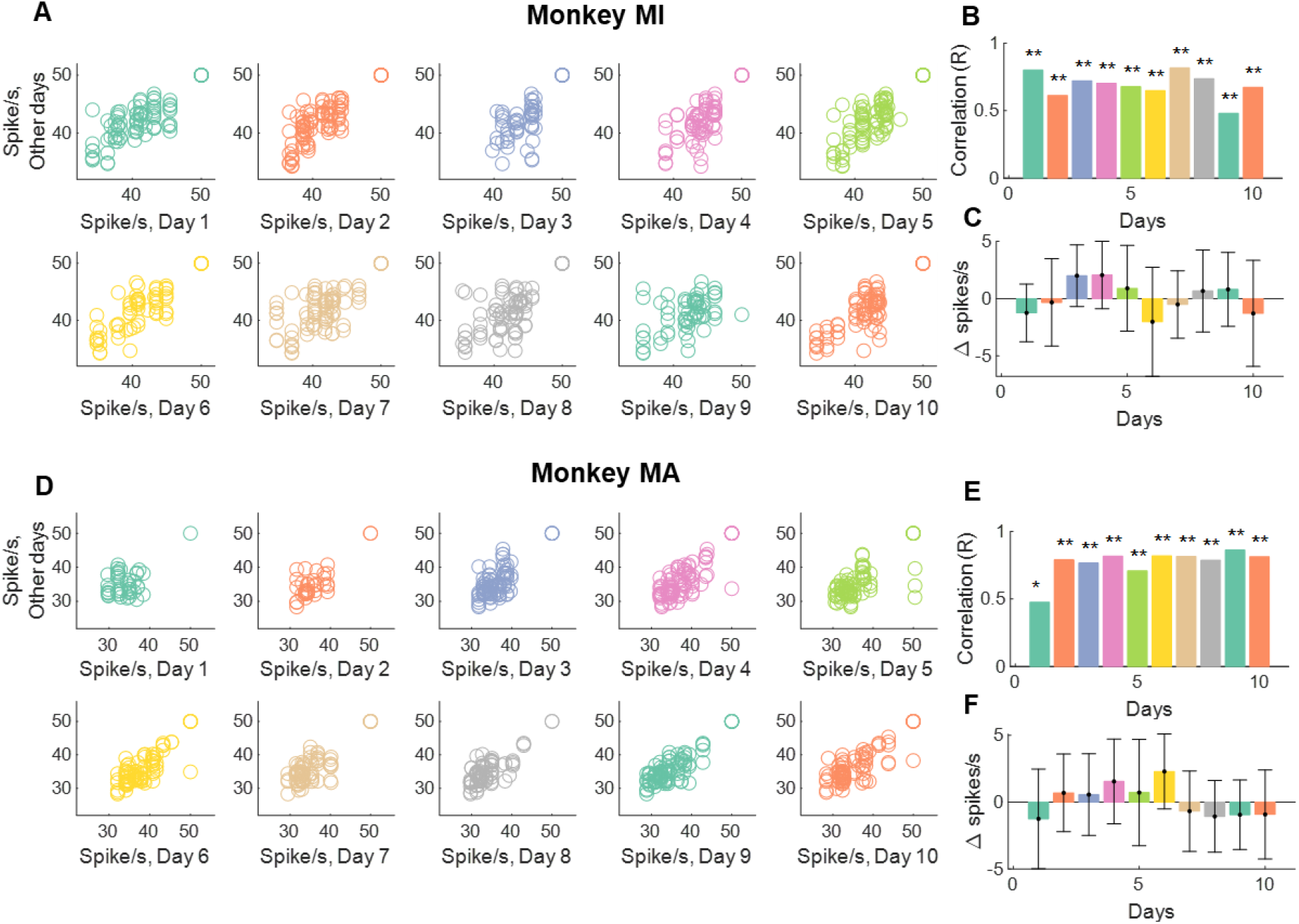
Motor unit discharge characteristics for the tracked motor units. **A.** The average instantaneous motor unit discharge rate was plotted for all tracked motor units at any given day. Note that some motor units may show different discharge rates because of changes in synaptic input. The day-to-day variability was very low (< 6 %) and this low variability is demonstrated by very high correlation values (**B**) for the tracked motor units. **C**. The absolute variability in discharge rate of the tracked motor units (i.e., the average motor unit discharge rate at day 1 minus the discharge rate of the same motor unit in the other days). Note that this correlation can only be significant if the motor units are tracked successfully, since the motor unit discharge rate shows high variability across the different units (see the figures below). **D-F.** The same plots as in **A-C** for Monkey (MA). *P<0.01, **P<0.001

**Figure 4.**
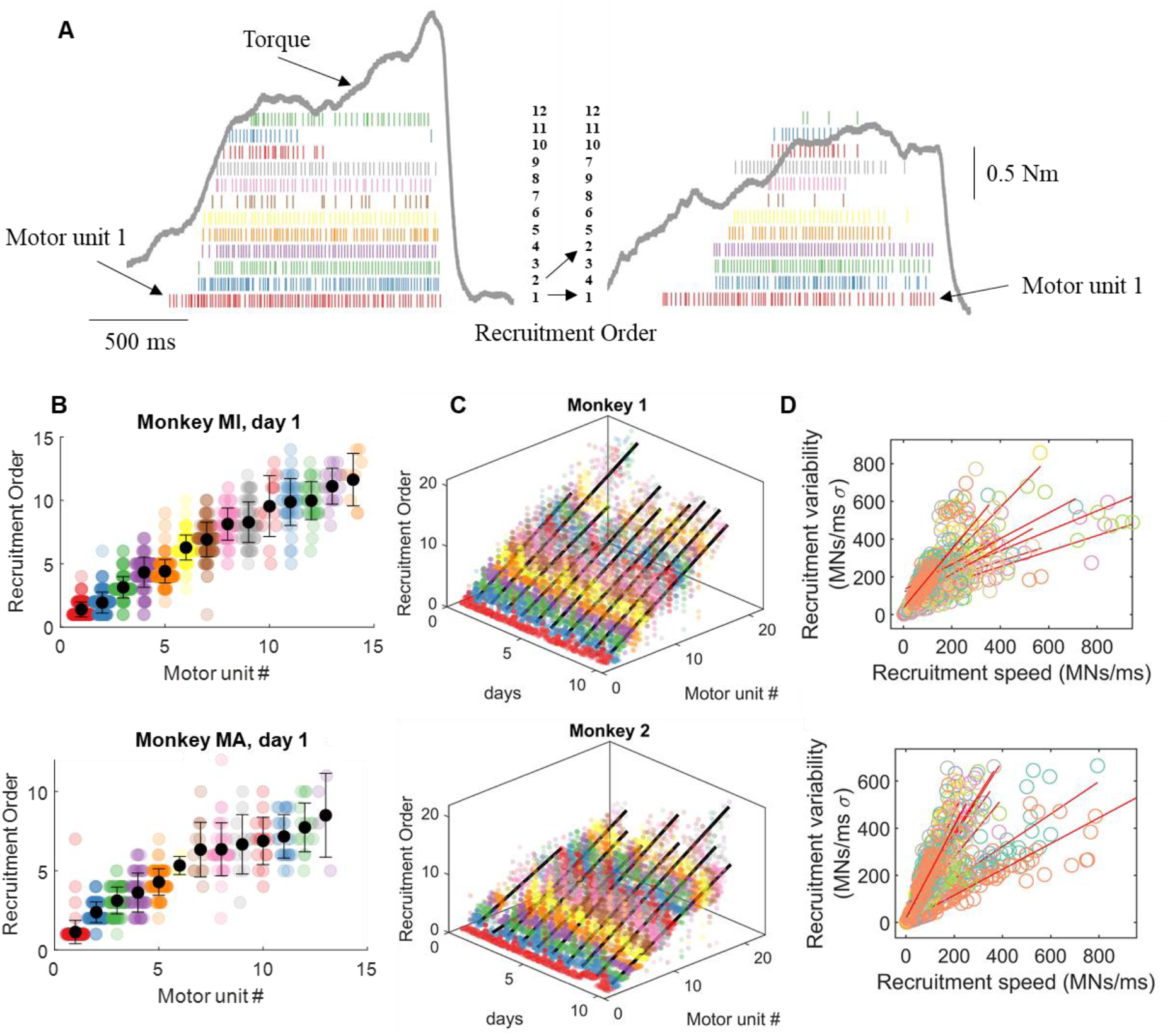
Motor unit recruitment thresholds and intervals across different contractions, motor units, and days. **A.** For each motor unit we calculated the shifts in recruitment order with respect to the average recruitment threshold of that unit. The motor units in the two contractions in **A** are color-coded with respect to the recruitment threshold in the first contraction. For example, it is possible to observe the shift in the recruitment order of #2 to #4 in the second contraction. However, these changes only happen for motor units with very similar threshold. For example, the motor unit red (#1 in left panel) and the highest threshold motor unit (#12 green) shows a consistent recruitment order. This can be well appreciated in the following figures when showing the motor unit recruitment order with respect to the average across the specific day. **B.** Swarm-plots of the recruitment order across all motor units. We first computed the recruitment threshold as the first spike of the motor unit during a specific contraction. We then averaged the recruitment threshold across all contractions for the specific motor unit that was tracked across all contractions (each dot in the swarm plot represents the recruitment threshold of a motor unit in an individual contraction). The average recruitment threshold was then used to sort the recruitment interval of all motor units. Note that each motor unit shows a stable behaviour across all contractions. **C.** Three-dimensional swarm plot for all the motor units across the 10 days. For both monkeys the relationship between recruitment order and motor unit number was linear across the 10 experimental sessions spaced over a month (R = 0.88 ± 0.04 for monkey MI and R = 0.88 ± 0.04 for monkey MA, P < 0.00001). **D.** The variability in recruitment order across days and contractions was highly correlated with the recruitment speed of motoneurons. The recruitment speed of motoneurons is an estimate of supraspinal drive and corresponds to the time derivative of the first discharge timings of all motor units during an individual contraction. Each regression line in D shows the variability across contractions for a specific day. Note the high variability in recruitment speed, which indicates the variance in rate of force development across the contractions for a specific day.

Previous evidence showed that motoneurons are recruited according to the size principle (*5*). This implies that for a given synaptic input, motoneurons are recruited according to intrinsic properties (*2*). However, some current and previous studies suggests a flexible control of spinal motor units in the mammalian nervous system (*7, 8*), so that a strict recruitment order is seen as a special case of a flexible control. According to this view, it is conceivable that variability in recruitment may occur over multiple experimental sessions where the monkeys are instructed to reach a target force level according to a broad range of contraction speeds. Contrary to this idea, we found a consistent recruitment order of motor units that was maintained across contractions and days (Figure 5). The recruitment order across the 10 experimental sessions was occasionally violated for motor units with very close recruitment thresholds (Fig. 5A-D). In these cases, the occasional reversals of recruitment order were highly correlated with the speed of recruitment (and therefore with the rate of force development) (Fig. 5C). With very fast recruitment, the difference in threshold between motor units with close recruitment threshold compresses to very small values so that the variability in synaptic input may likely explain the occasional reversals (that happened in a small range).

**Figure 5.**
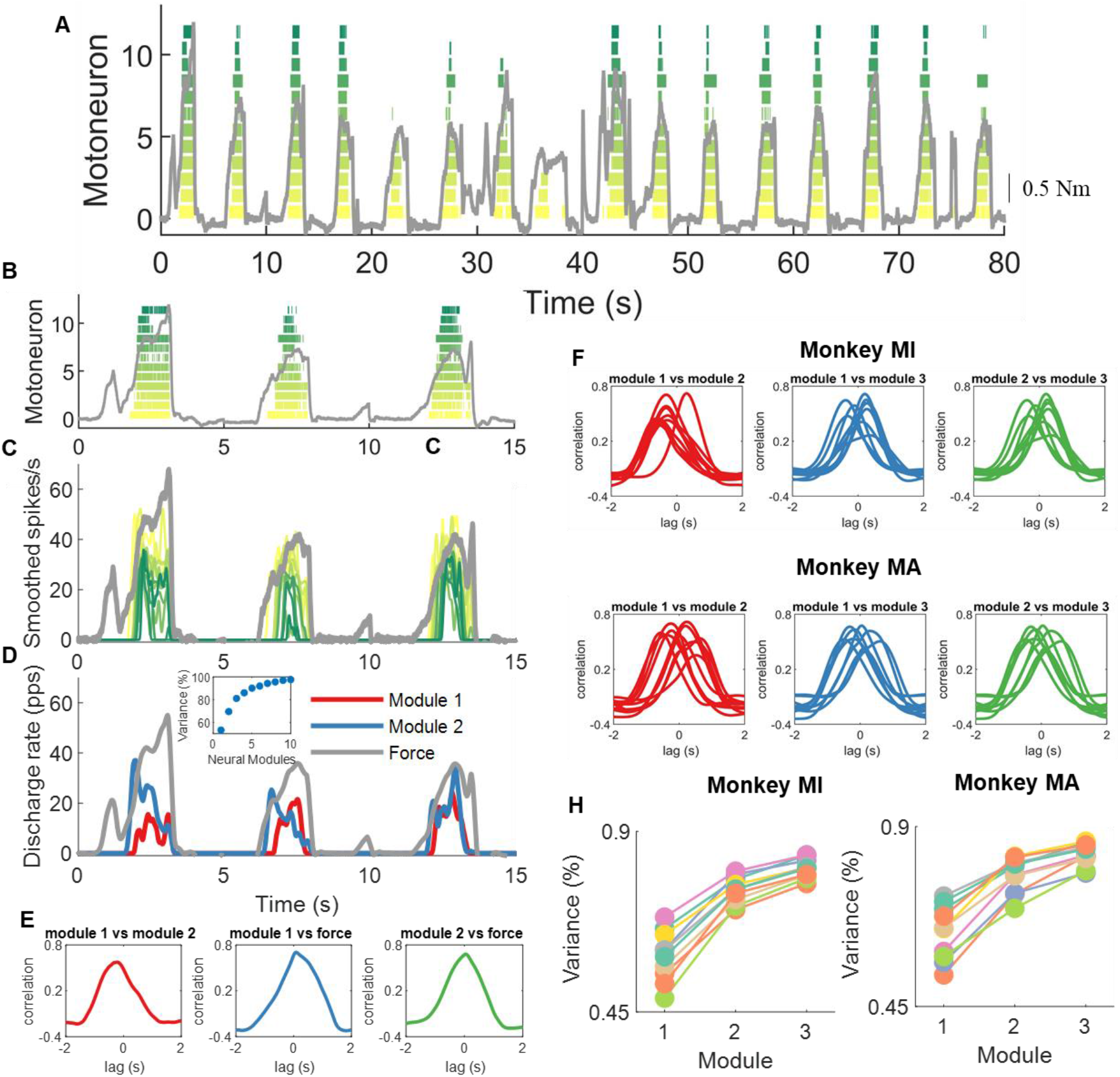
Encoding of muscle force by motor units. We aimed at decoding and encoding the temporal motor unit information into components by non-negative matrix factorization. **A**. Raster plot of twelve motor units during a subset of macaque voluntary isometric contractions (grey lines indicate the torque signal). Note the variability in peak forces and rate of force developments. **B-C-D** show the first three contractions in A. **C.** The motor unit spike trains in A were convoluted with a 2.5 Hz Hanning window. Note the high correlation between the motor unit smoothed discharge rates and muscle force. **D.** We applied the reduction dimensionality technique non-negative matrix factorization. We constrained the model to learn the components in the motor unit discharge rates up to 10 factors. In this example, the two modules that together explained approximately 80% of the variance are shown. Note that these two modules are highly correlated, and time shifted. The inset in D shows the reconstruction accuracy (variance %) of the neural modules with respect to the original signal (smoothed motor unit discharge rates). **E.** We applied cross-correlation analysis between the modules and muscle force. This example shows the correlation between the first module and second as well with voluntary force. The same method was then applied for all modules in both monkeys, which are shown in **F.** Note the high correlation across all days and for both monkeys. Moreover, there was always one module with a dominant component (the lag between the different modules was never zero). This indicates that there is only one component constrained by the size principle, since the motor unit recruitment thresholds are highly preserved across all contractions **H**. The reconstruction accuracy (variance %) explained across the 10 days for both monkeys.

The present results are in accordance with previous human and in-vitro experiments indicating that motor units are recruited in a specific order. We therefore wanted to understand if there are specific patterns in the motor unit discharge timings that control the recruitment and muscle force. We applied a non-negative matrix factorization analysis (*22*) to the motor unit discharge timings. Because of the large amount of motor unit data, we were able to discern the exact patterns common to all and to sub-groups of motor units.

The non-negative matrix factorization revealed a principal component that explained ~50% of the variance. This component was present in the activity of virtually all low-thresholds motor units. There was a significant second factor that explained ~25% of the variance. Interestingly, this originated mainly from high threshold motor units and was an undistorted, time-shifted version of the first component. We then performed correlation analysis between all the components (10 in total, see Methods) and looked at the specific weight distributions across the individual motor unit recruitment thresholds. We found that these components were consistently time-shifted and with very high correlation values between each other (Figure 5F). Moreover, the second component was consistently present only in the high-threshold motor units. These results indicate that motor unit discharge rates during natural tasks in macaque monkeys are driven by one dominant command, which manifests in time-shifted form because of the progressive recruitment imposed by the size principle. Because the motoneuron is a non-linear system, the ensemble activity strongly indicates that these common fluctuations must originate from common input from cortical, afferents, or brainstem pathways. We provide strong evidence that a main component drives a pool of macaque brachioradialis motor units that is mediated by the recruitment order of the motor units.

### Motor unit synchronization

It has been reported that the discharge timings of spinal motor units show very high synchronization values (*2*), which are associated with the generation of muscle force (*23*). Accordingly, we found high values of motor unit synchronization similar to what is typically observed in humans (*24*). We analysed synchronization in two frequency bandwidths; one which retains most of the information of the corticospinal pathways, 0-40 Hz (*25*), and a narrowed one (0-5 Hz), which retains the information that is correlated to force generation (corresponding to the muscle low-pass filtering bandwidth, <5Hz (*26*)). The cross-correlation value for the low pass filtered signals (5 Hz) at lag 0 was 0.78 ± 0.01 and 0.72 ± 0.10 for MI and MA, respectively. The values across the different bandwidths were consistently very high. These values also showed very small deviations across the contractions (1.55% and 2.30% for MI and MA; see Figure 6). Interestingly, the value of synchronization was in the highest portion of the range observed in humans (R = 0.5 – 0.8).

**Figure 6.**
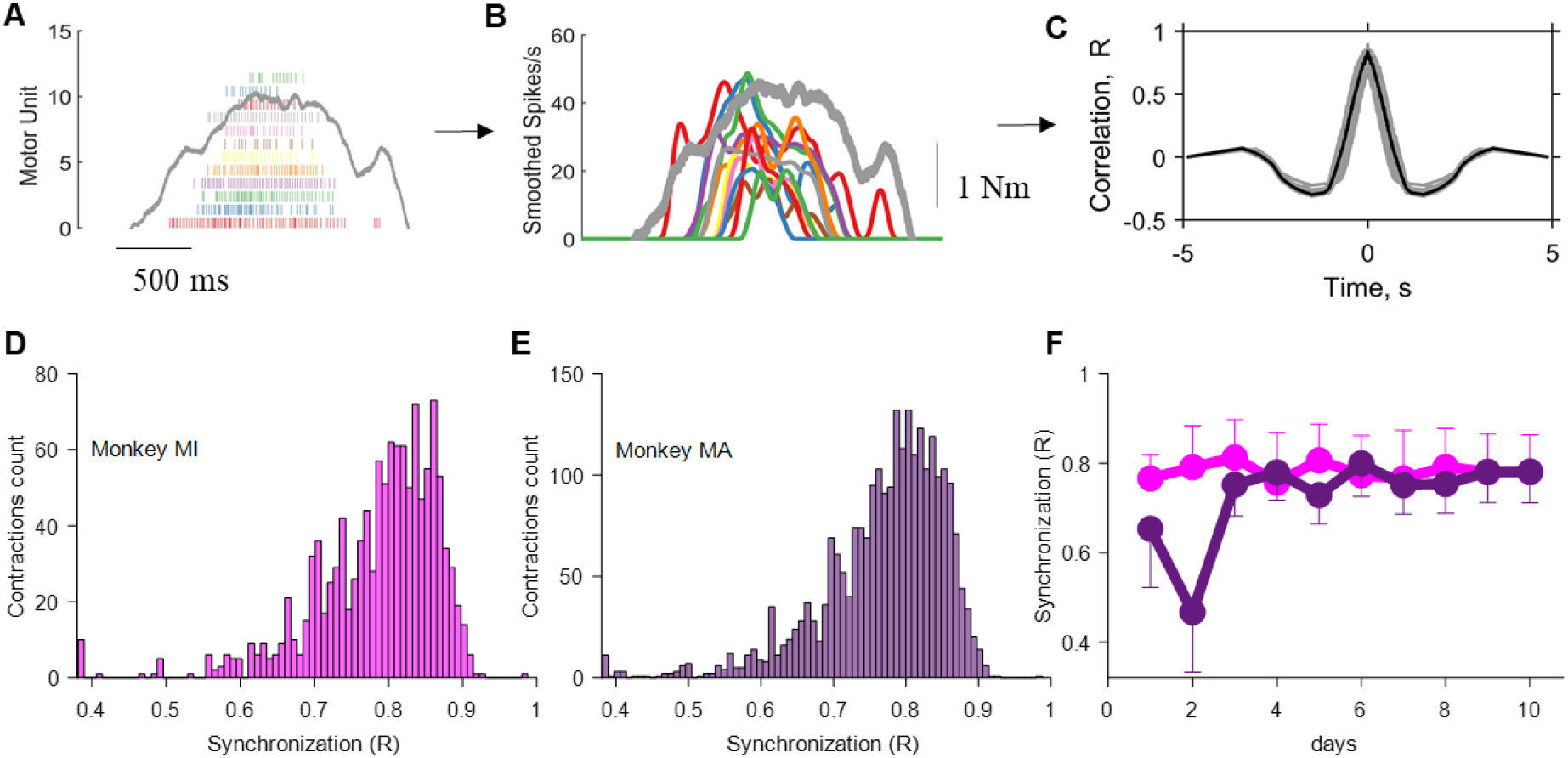
Motor unit synchronization across the different contractions for monkey MI and MA (color-coded). **A-C** Pipeline for the estimate of motor unit synchronization for an individual contraction. **A.** Raster plot of 12 motor units (grey indicates the torque with scale shown in panel B). **B.** The discharge timings of the motor units were filtered with a Hanning window of 200 ms. **C**. The synchronization value was obtained by performing the cross-correlation function between two groups of randomly permutated groups of motor units (number of permutations = 100). Note that the synchronization value was relatively high, and comparable to what observed in humans during rapid force contractions. **D-E.** Histogram of the synchronization value across the individual contractions for both monkeys. **F.** The synchronization value was stable across the ten days (average and standard deviation for each days are shown). For Monkey 2 the first two days resulted in a lower synchronization value due to a lower number of identified motor units, as shown previously. Note that the small variability in synchronization value in D and E was fully explained by the instantaneous discharge rate of the motor units, as previously shown (*24, 27*).

The high correlation further indicates that the motoneurons likely received a strong common excitatory synaptic input and that this input was stable across days (Fig. 6F).

### Variability of motor commands are distributed within and between motor unit pools and have a common supraspinal origin

The previous results indicated that despite a large range of values in rate of force development and motor unit recruitment discharge characteristics, the general motor control scheme shows high reliability in the recruitment order and neural output of brachioradialis motor units. We also monitored the activity of other muscles involved in the tasks to verify behavioural variability across trials and days. We implanted 16 intramuscular EMG (iEMG) electrodes into the muscles of the left and right arm (Figure 7) and nerve cuffs around the median and radial nerves. The recordings from the iEMG signals were performed for the voluntary force contractions as well as for the involuntary stimulated contractions. We investigated the full bandwidth of efferent and afferent volleys with small changes of electric currents applied on the axon, until maximum efferent activation (M-wave).

**Figure 7.**
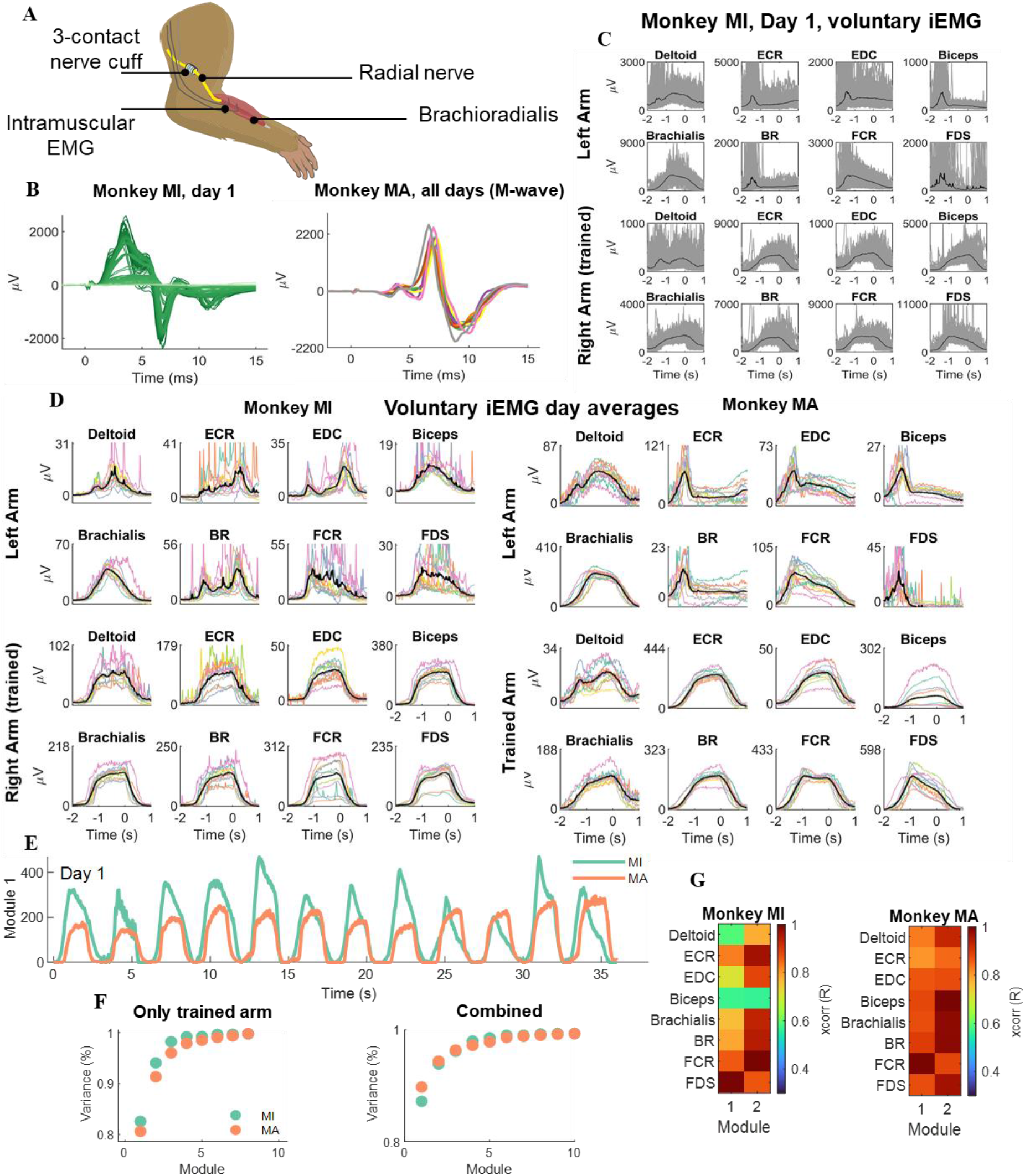
Neuromuscular implants in macaques. **A**. Both monkeys were implanted bilaterally with a nerve cuff around the median and radial nerves. Implanted intramuscular EMG signals recorded the gross myoelectric activity of 16 muscles bilaterally (8 muscles per side). **B.** During each experiment, the nerve cuff delivered stimulation pulses at supramaximal intensity (M-waves) and ramped down in small decrements of 0.1μA. The left side of panel B (dark green lines) shows the iEMG recording sessions from supramaximal intensity to the smallest (light green). On the right side of the panel twelve M-waves obtained during the different days (color-coded). **C.** The iEMG signals from the voluntary contractions during one experimental session. Individual contractions as well as the average (black line) are shown. Note the high intertrial variability in gross EMG responses. **D.** The average iEMG traces across days (color-coded), for monkey MI and MA. **E.** Non-negative matrix factorization analysis applied to the gross iEMG signals. The neural modules that explained most of the variance are shown for each monkey. **F.** The reconstruction accuracy (variance %) of the components extracted by NNMF. Note that one component explained more than 80% of the variance. **G.** The cross-correlation of the first two modules for the respective muscles.

The potentials evoked by electrical stimulation showed high reliability across days, with negligible deviations around the mean (Figure 7). This demonstrated stability of the recordings over days. On the other hand, the voluntary EMG amplitudes showed very high variability, with some muscles (including the brachioradialis) showing a 2-fold difference in maximal amplitude. This indicated relatively large variability in the way contractions were executed.

We then applied the same method for the identification of motor unit components (Fig. 5) to identify the neural modules within the muscles, as classically referred to as muscle synergies (*28–30*). We found one invariant neural component that explained more than 90% of the variance. This component was present either in the iEMG signals only from the trained limb, or in the combined iEMG signals from both limbs. This result further supports the role of a common input that is distributed between and within motor nuclei that is processed by the size principle and spinal cord circuitries, despite the large variability in the muscle activities.

## DISCUSSION

We have proposed a new non-invasive method based on wearable sensors to monitor spinal motoneurons in non-human primates that surpasses previous invasive methods in terms of performance (number of motor units), accuracy, and the possibility to track units over time. With this method, we reveal an accurate representation of the strategies used by the nervous system to control motor units and muscle force. The condensed spatial dimensions given by the high-density grids allowed us to identify the same motor units in two macaque monkeys performing natural isometric contractions across several experimental sessions. The access to populations of spinal motor units and their longitudinal tracking provides a framework to study the changes in recruitment of spinal motoneurons and rate coding during natural tasks.

With respect to intramuscular recordings, these non-invasive approaches provide stable signals even during fast contractions (*31*), a greater number of decoded motor units (*21*), and the possibility to track the same motor units over multiple experimental sessions across days (*32*) and weeks (*20*). These approaches have been developed and extensively validated in humans (*20, 21, 33*). Here, for the first time, we show a non-invasive framework for decoding and longitudinally tracking relatively large populations of spinal motor neurons in behaving monkeys.

We found relatively high motor unit discharge rates in macaque monkeys (41.7 ± 1.4, 38.4 ± 2.0 spikes/s for MI and MA respectively across the ten days). These discharge rates were higher than those observed in isometric contractions at low and moderate forces in humans (<50 % of maximal voluntary force, <30 spikes/s) (*33*). Conversely, when related to fast human isometric contractions of the tibialis anterior muscle, the observed rates are similar (40.09 and 42.85 spikes/s, for the non-human and human motor units, respectively (*31*)).

The discharge timings of the motor units represent the neural code that generates muscle force. Recordings of motor unit activity during voluntary force contractions allow us to test the recruitment of motor units by the central nervous system in a detailed way, clarifying current debates in motor control. It has been debated for decades whether the common motoneuron fluctuations observed at the motor unit level are an epiphenomenon or have a functional origin. Similarly, the Henneman size principle has been constantly under investigation, due to the lack of in-vivo evidence with contractions at different rates of force development (*6-9, 34, 35*). These problems arise because of the lack of adequate methods.

Here we showed that the neural drive to the muscle is highly structured in a hierarchical fashion. We found strong associations between hierarchy and behaviour, so that for a given common input signal, the motoneurons behave synchronously once they reach their threshold to discharge, likely dictated by the intrinsic motoneuron properties. Our results are in strong accordance with simulations suggesting that the spinal cord decodes inputs from descending pathways by modulating the recruitment and derecruitment of motoneurons (*36*). The factorization analysis applied to individual motor unit discharge timings and gross intramuscular EMG signals from the trained and untrained limb, revealed that one component explained more than 80% of the variance. The motor unit findings revealed that this component is filtered by size principle. Our results demonstrate the interplay between common synaptic input and size principle.

In conclusion we presented a new non-invasive framework to decode populations of single spinal neural cells in macaque monkeys, which allows us to move from simple measures of behaviour (force) to the inputs that determine that behaviour. In addition to being non-invasive, this framework identifies the same motor units across months over the full force range. This is critical since inferring the patterns of motor behaviour by random sampling small population of active units may be inadequate (*7, 11, 16, 17, 37*). We anticipate that this approach may find further utility when combined with invasive recordings of central motor circuits, which can provide direct access to the various putative sources of common drive (*38–40*).

## Materials and Methods

### Animals

Recordings were performed from two adult female awake behaving monkeys (*M. mulatta*; monkeys MI and MA, age 6, weight 6.2 and 6.7 kg respectively). All animal procedures were performed under appropriate licences issued by the UK Home Office in accordance with the Animals (Scientific Procedures) Act (1986) and were approved by the Animal Welfare and Ethical Review Board of Newcastle University.

### Behavioural Task

The monkeys were trained to perform an isometric elbow flexion task with their right arm. Monkey MA was also trained to perform this task with her left arm. The forearm was placed into a rigid plastic cast. This was 3D printed from a digital model of the forearm made using a laser scanner (Go!Scan, Creaform 3D, Levis, Quebec, Canada), ensuring a close but comfortable fit. A further support held the upper arm; the supports were attached to the training cage to fix the elbow in 90° flexion, and the forearm in semi-pronation so that the radius and ulnar were oriented in a vertical plane. A load cell (LC703-25; OMEGA Engineering Inc., Norwalk, CT, USA) attached to the forearm cast registered elbow flexion torque. The force (kgF) applied to the load cell was recorded as a voltage signal by a custom designed task programme. A calibration factor was determined which allowed for the conversion of the voltage signal back into kilogram force (kgF) at a later stage. To determine the torque (N·m) produced by the animals, the recorded kilogram force was gravity corrected and converted into Newtons (N) and secondly multiplied by the distance between the load cell sensor and the elbow pivot joint (0.08m). The monkey initiated a trial by contracting elbow flexors to place the torque within a set window (1.648-3.295 N·m). This window was kept constant in all sessions and for both animals. The torque had to be held in this window for 1 s before releasing to obtain a food reward. Auditory cues were used to indicate to the monkey that the exerted force was within the required window, or else it was too high. Auditory feedback was also given to mark the end of the hold period. Recordings were collected from 10 sessions spanning 30 and 24 days for monkey MI and MA, respectively.

### Surgical Preparation

After behavioural training was complete, monkey MI underwent a sterile implant surgery. After initial sedation with ketamine (10mg·kg^-1^ IM), anaesthesia was induced with medetomidine (3 μg·kg^-1^ IM) and midazolam (0.3mg·kg^-1^ IM). The animal was then intubated and anaesthesia maintained using inhalation of sevoflurane (2.5-3.5% in 100% O_2_) and IV infusion of alfentanil (0.4 μg·kg^-1^·min^-1^). Methylprednisolone was infused to reduce oedema (5.4mg·kg^-1^·hr^-1^ IV). Blood-oxygen saturation, heart rate, arterial blood pressure (using a non-invasive blood pressure cuff on the leg), core and peripheral temperature and end-tidal CO_2_ were monitored throughout; ventilation was supported with a positive pressure ventilator. Hartmann’s solution was infused to prevent dehydration (total infusion rate including drug solutions 5-10 ml·kg^-1^·h^-1^). Body temperature was maintained at 37°C using a thermostatically controlled heating blanket and also a source of warmed air. Intraoperative prophylactic antibiotics (cefotaxime 20mg·kg^-1^ IV) and analgesia (carprofen 5 mg·kg^-1^ SC) were given.

In monkey MI, nerve cuff electrodes (Microprobe, Gaithersburg, MD, USA) were implanted around the median and deep radial nerves bilaterally and secured with the integral sutures. Each cuff contained eight contacts, arranged as two sets of four wires placed radially around the inner circumference. A plastic headpiece (TECAPEEK MT CF30, Ensinger, Nufringen, Germany) was manufactured based on an MRI scan to fit the skull and fixed using ceramic bone screws (Thomas Recording Inc, Giessen, Germany) and dental acrylic. Intramuscular electrodes comprising Teflon-insulated stainless-steel wires were implanted in eight arm and forearm muscles bilaterally for gross electromyography (EMG) recording. Specifically, the muscles that were implanted with intramuscular electrodes corresponded to: deltoids, extensor carpi radialis (ECR), extensor digitorum communis (EDC), biceps brachii, brachialis, brachioradialis, flexor carpi radialis, and the flexor digitorum superficialis muscle (FD). The EMG and nerve cuff wires were tunnelled subcutaneously to connectors fixed to the headpiece. Nine weeks after monkey MI’s first implant surgery, several wires connected to the deep radial nerve cuffs bilaterally were found to be broken, and stimulation through these cuffs was no longer possible. Replacement cuffs (with three contacts each, organised radially around the inner circumference) were then implanted bilaterally on the radial nerve below the spiral groove in a further brief surgery, again with wires tunnelled subcutaneously to the head. Monkey MA underwent the implant surgery at a later stage to monkey MI and so was implanted with the same three contact cuffs around the median and radial nerves, along with EMG electrodes in the same muscles and fitted headpiece. All recordings were subsequently collected using the three contact nerve cuffs.

Post-operative care included a full programme of antibiotic (co-amoxiclav, dose as above) and analgesics (meloxicam, 0.2mg kg^-1^ oral plus a single dose of buprenorphine 0.02mg kg^-1^ IM).

### Nerve cuff stimulation and recording

Biopolar current pulses (0.2ms per phase) were delivered through the first and third contacts of the three contact radial cuffs with a bi-phasic constant current isolated stimulator (Model DS4, Digitimer, Hertfordshire, UK). Stimulus current was delivered at supramaximal intensity (0.45mA for monkey MI and 0.4mA for monkey MA) and ramped down in decrements of 0.1μA to threshold intensity. Left and right arms were stimulated in different sessions, following recordings of the motor task.

### Electrophysiological Recordings

Recordings were made from the brachioradialis muscle using a high-density surface EMG grids (GR04MMI305, OT Bioelettronica, Turin, Italy) with 64 electrodes (spacing 4mm). A bi-adhesive foam strip with holes aligned to the matrix was placed on the grid, and the holes filled with conductive paste (CC1, OT Bioelettronica, Turin, Italy). This assembly was then stuck to the skin over the muscle. To ensure good skin contact the forearm was shaved and cleansed with alcohol wipes. The location of the grid on the skin was marked each day with permanent marker pen to ensure reproducible placement from session to session. Standard surface adhesive electrodes (Neuroline 720; Ambu A/S, Ballerup, Denmark) were placed over the flexor and extensor tendons at the wrist to act as reference and ground; in the implanted animal (monkey MI), one of the unused nerve cuff electrodes was used as the ground. The surface grid electrode was connected to a custom printed circuit board containing a 64-channel amplifier (gain 192; bandwidth 30Hz - 2 kHz) and an analogue-to-digital convertor (RHD2164; Intan Technologies LLC, Los Angeles, CA, USA). Digitized signals were sent over a serial peripheral interface (SPI) cable to an RHD USB interface board (also Intan Technologies). This allowed data to be captured to a computer hard disc (5 kSamples/s) along with the elbow torque signal and digital markers signalling the phases of task performance and stimulus timing. Voluntary brachioradialis activity was recorded from the grid electrode during performance of the behavioural task (typically 100 successful trials per session). Involuntary contractions were recorded by the intramuscular electrodes and the grid electrode during the radial nerve stimulation protocol.

### Motor unit decomposition and analysis

The high-density EMG recordings were offline digitally filtered with a 20-500 Hz Butterworth filter. Semi-automated MATLAB software extracted the area under the power spectrum and amplitude of each of the 64 channels and highlighted the channels with poor signal to noise ratio for visual inspection and exclusion from subsequent analysis. After this procedure, the monopolar signals were used for the decomposition. Identification of the individual motor unit firings was accomplished through a previously proposed algorithm (*21*), modified for these large datasets to use a graphical processing unit (GPU) running CUDA software (Nvidia Inc, Santa Clara, California, USA).

Briefly, this algorithm takes advantage of the unique two-dimensional spatiotemporal features of individual motor unit action potentials, to converge on an estimate of the motor unit spike trains. The decomposition blindly identifies the motor unit firings; only motor units with high silhouette-measure (>0.92 SIL) are initially maintained. SIL represents a qualitative measure of decomposition accuracy which is comparable to the pulse to noise ratio, ranging from 0 to 1, where 1 indicates perfect clustering of the motor unit action potential. The blind source separation procedure leverages the high spatial and temporal dimensionality of motor unit action potentials. This information is used to converge in an iterative way in the unique time-series representation of the firing times of the alpha motoneurons. We briefly describe here the general steps of decomposition. For a more detailed look into the details of high density EMG decomposition, the technical and physiological details have been described previously (*41, 42*)

The EMG signal corresponds to the filtering of the motoneuron action potential by the muscle tissue with some added noise. Therefore, it is possible to represent in a mathematical form the signal that is carried by each channel of a multidimensional arrays of EMG signals. The EMG signal can be described as a convolution of the motoneuron discharge timings (sources) by the muscle tissue (muscle unit action potentials). The sources (s) are the motoneuron axonal action potentials when reaching the muscle fibres and can be written as Dirac delta function.

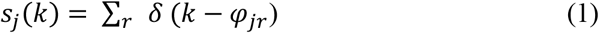

where *φ_jr_* represent the spike times of the *j*th motor unit. We can then write the EMG signal in a matrix 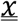 form (e.g., when recorded with multidimensional arrays such as the high-density EMG grids used in this study) as:

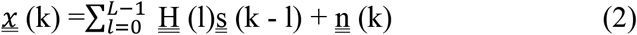

where 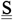 (k) = [s_1_ (k), s_2_ (k),…, S_n_ (k)]^T^ represent the n motor unit discharge times that generate the EMG signal 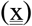 and 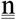 is the noise to for each electrode. The matrix 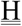 (l) in eq. 2 contains the spatial information of the motor unit action potential and has size m *x* l with l*th* sample of motor unit action potentials for the *n* motor units and *m* channels (two-dimensional format, hereafter referred to 2D motor unit waveform). The high spatial sampling given by the 64 electrodes further enhanced by extending the observation numbers (*41*) allows the recovery of the sources in an iterative blind way with a function that maximizes the sparsity between each motor unit action potential (Fig. 1A). This process is obtained in a fully automatic and blind way; therefore, we can inspect the validity of decomposition by spike-triggered averaging. With spike-trigger averaging it is also possible to retrieve by correlation analysis the information that is carried by the action potential 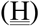 in different days, in a fully automatic way. By using 2D correlation analysis it indeed possible track motor unit waveform across weeks (*32*) and even months (*24*). The motor unit tracking uses the information carried in 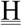 to compare across sessions the two dimensional cross-correlations across all possible combinations of motor unit action potentials. The two-dimensional cross-correlation (2D correlation hereafter) is comparable to a one-dimensional cross-correlation, but with a weighted average across the time-space features of the motor unit waveforms (see Figure 1). The output of the two-dimensional cross-correlation ranges from 0 to 1, where 1 indicate maximal similarity. For example, two randomly-selected motor units have a two-dimensional cross-correlation lower than 0.3 (*20*).

### Motor unit characteristics

We first displayed all motor units 2D correlation values with R > 0.55 and with a total number of discharge timings (impulses) >100 and visually inspected the waveforms for potential errors. The unique combinations of motor unit waveform that were preserved after this visual inspection stage had R > 0.70. From all the retained motor units, we computed the instantaneous discharge rate (inverse of the inter-spike interval) averaged across the hold period for all trials of the task on a given day. Synchronization of the motor unit pool was also assessed, as the magnitude of the cross-correlation between two equally sized groups of motor unit spike trains. The number of motor units in each group was randomly assigned for a total of 100 permutations. For each iteration two random unique subsets of units were selected for each group (each group being half of the total number of the identified units during a specific contraction). The spike trains (binary signals) for each motor unit group were then summed and smoothed using a Hann window with two corner frequencies of 40 Hz and 2.5 Hz. We chose two Hann window because this cut-off retains most of the oscillatory activity of the motoneuron pool (40 Hz) and the low frequency is mainly associated to the neural drive that is responsible for force production (i.e., the correlation between a force signal and the low pass filtered motor unit discharge timings is minimally distorted by the musculotendinous unit).

For the motor unit recruitment threshold estimates, we first looked at the recruitment order (in seconds) of the motor unit during the individual contractions. This was estimated by taking the time point when that unit was active for the first time. We then calculated the average recruitment threshold (in seconds) for all units across all contractions. Afterwards, we labelled each unit from 1 to the maximum number of identified units in a specific contraction (i.e., a motor unit takes the value of 1 if it is the first recruited). Then we plotted the recruitment thresholds for each specific unit across all contractions. Because the labelling is not dependent on the average, if there is a correlation between the average recruitment threshold and the binarized recruitment threshold across all contractions, this relationship indicates the amount of flexibility in recruitment order obtained by the nervous system in a direct way.

We then computed the derivative of the recruitment of the motor units. After calculating the recruitment thresholds (in seconds) of all the motor units in each contraction, the recruitment threshold was sorted from the smallest to the largest and we computed the derivative of this vector. The derivative of this vector corresponds to the number of motor units recruited per seconds, which is an estimate of the efferent drive received by the population of motor units, i.e., a faster recruitment speed of motoneurons results in a faster rate of force development (*31, 43*). We then associated for each contraction the variability in recruitment order, that was calculated as the standard deviation of the binarized recruitment thresholds versus the motor unit recruitment speed (the first derivative of the recruitment thresholds). If there would be an association between these two variables it would indicate that a faster recruitment (which could be due to higher synaptic input) is associated with a violation in the recruitment order.

### Factorization of motor unit activities

We factorized the motor unit discharge timings with a non-negative matrix factorization method (NNMF, *29*). This method can learn specific features in 2D images such human face characteristics or sematic properties of a written text with the use of linear algebra. In the context of neural signals, we constrained this method to learn the unique components in the motor unit discharge rates that are responsible for force production. Figure 6 shows the overall architecture for this analysis.

The force level developed by a muscle is driven by the number of motor unit activation signals, which can be represented as time sequences of M dimensional vectors, that correspond to the activation of the motoneurons **m(t)** in response to common and independent synaptic inputs arising from afferent and efferent volleys. Therefore, we can express the motoneuron behaviour as combinations of N varying synaptic inputs which construct a specific motor unit firing characteristic, or *neural module*, expressed as {*w_i_*(*t*)}_*i=1,…N*_

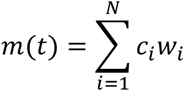

where *c_i_* is a non-negative scaling coefficient of the *i*-th neural module. We are interested in finding the *W_i_* vectors within the low-frequency motor unit discharge rates. Because motor unit firing rates are non-negative, we can utilize NNMF (*22*) to constrain *w_i_* to be non-negative. This procedure maximizes the interpretability of the data since the representation of the neural motor unit ensemble only includes additive and not subtractive combinations, therefore having an output module with the same scale as the input signal. NNMF iteratively finds the non-negative factors W and H with an interactive procedure that minimize the residuals between **D** (the sources) and **W*H**. So that W*H is a lower-rank approximation of the firings of the individual motor units (D). The firing of the individual motor units are stored in a matrix with rows equal to the number of identified motor units and with columns having the duration of the recording. The motor unit are initially stored as Dirac delta’s function *δ*(*k* – *φ_jr_*), and then low pass filtered at 2.5 Hz (Figure 6). NNMF is an iterative algorithm that starts with random initial value of W and H. Because the root mean square of D can have local minima, we performed up to 1000 iterations to converge to a representative reconstruction of D = W*H.

We then evaluated the output of NNMF with different decoding-encoding functions. First, we constrained the number of factors number equal to the number of identified motor units across a specific day. After this initial procedure, we consistently found that >10 factor explained 99% of the variance. The reconstruction accuracy (residual variance or variance explained) was calculated by computing the residuals (D - W*H) and then computing the deviation from the mean (R^2^). Second, we evaluated the decomposition by looking at the decoding-encoding of the individual neurons with the respect to the matrix W. This analysis was computed by performing the cross-correlation between the low-pass filtered motor unit discharge rates (D) and the individual neural modules (W) extracted by NNMF. The same method was applied on the gross EMG signals from the intramuscular electrode. After rectification and averaging, the average EMG signals for each day were processed by NNMF and the residual variance was calculated in the same way for the motor units (Figure 7 shows the results and analysis of the intramuscular EMG signals). All of the analyses were performed in MATLAB.

## Funding

Supported by a grant from The William Leech Charity. JI received the support from “la Caixa” Foundation (ID 100010434; fellowship code LCF/BQ/PI21/11830018).

## Notes

### Competing Interest Statement

The authors have declared no competing interest.

